# Rapid *in vitro* activity of telavancin against *Bacillus anthracis* and *in vivo* protection against inhalation anthrax infection in the rabbit model

**DOI:** 10.1101/2023.02.13.528259

**Authors:** William S. Lawrence, Jennifer E. Peel, Richard Slayden, Johnny W. Peterson, Wallace B. Baze, Martha E. Hensel, Elbert B. Whorton, David W.C. Beasley, Jason E. Cummings, Ines Macias-Perez

## Abstract

Anthrax, caused by the bacterium *Bacillus anthracis*, is a zoonotic disease that manifests in various forms in human infection, depending on the route of infection. Inhalation anthrax, the most detrimental form of the disease, comes about from the inhalation of anthrax spores and progresses to severe life-threatening conditions late in infection. Notably, there are FDA-approved antibiotics that are effective at treating the disease when administered promptly; however, these antibiotics would be rendered useless against strains of *B. anthracis* that were genetically modified to be resistant to these antibiotics. Consequently, the search for new and effective therapeutics to combat anthrax infection continues. In this study, telavancin (Vibativ^®^), a semisynthetic lipoglycopeptide antibiotic, was assessed for *in vitro* activity against 17 *B. anthracis* strains and tested for the protective efficacy against inhalation anthrax infection in the rabbit model. Telavancin demonstrated potent *in vitro* activity against *B. anthracis* which led us to test its efficacy in the rabbit inhalation anthrax model. Rabbits were infected with a lethal dose of anthrax spores via the inhalation route and treated intravenously with telavancin at 30 mg/kg every 12 hours, a dose that mimics the levels measured in the serum of humans, for 5 days upon detection of antigenemia. Blood samples were collected at various times post-infection to assess the level of bacteremia and antibody production, and tissues were collected to determine bacterial load. The animals’ body temperatures were also recorded. Telavancin conveyed 100% survival in this model. Moreover, the dosage of telavancin used for the study effectively cleared *B. anthracis* from the bloodstream and organ tissues, even more effectively than a humanized dose of levofloxacin. Collectively, the low MICs against all strains tested and rapid bactericidal *in vivo* activity demonstrate that telavancin has the potential to be an effective alternative for the treatment or prophylaxis of anthrax infection.

**Author Summary:** *Bacillus anthracis*, the causative agent of anthrax, continues to interest the research community due to its past and future potential use as bioweapon. Importantly, as a bacterial pathogen, *B. anthracis* is capable of developing resistance to the antibiotics currently used to treat the infection, either naturally or by deliberate, nefarious means. Consequently, there remains a need to discover, develop, and/or repurpose new antibiotics that would be effective at treating anthrax infection. In this study, we evaluated the antibacterial activity of telavancin, a semisynthetic glycopeptide antibiotic clinically approved to treat complicated skin and skin structure infections, against various strains of *B. anthracis in vitro*, and we assessed the protective efficacy of telavancin against inhalation anthrax infection in the rabbit model. We show that telavancin is very potent against numerous *B. anthracis* strains *in vitro*, and its level of potency surpassed that of another antibiotic currently approved and used to prevent anthrax infection. Moreover, we show that telavancin protects against inhalation anthrax infection *in vivo*. Overall, our findings support the use of telavancin as an effective therapeutic for anthrax infection.

## Introduction

*Bacillus anthracis* is a Gram-positive, spore-forming bacillus bacterium that causes anthrax infection. It is a category A bioterrorism agent because it is “…infective in low doses…suitable for mass production, storage, and weaponization; [and] stable during dissemination” [1]. Depending on the route of exposure, anthrax infection manifests in four forms: cutaneous, gastrointestinal, inhalational, and injectional. Inhalation anthrax infection is the most detrimental form of the disease, with a fatality rate of up to 80% when either no treatment is given, or treatment is delayed [2, 3]. The disease occurs from inhalation of anthrax spores which, due to their small size, can reach the alveolar spaces of the lungs, where macrophages and dendritic cells phagocytose them. These immune cells carrying the spores then migrate to regional lymph nodes where the anthrax spores germinate into vegetative bacteria and replicate. Eventually, the bacteria enter the lymphatic system and bloodstream resulting in systemic anthrax infection. Initial symptoms of the disease are nonspecific and include fever, chills, body aches, fatigue, and nausea. Still, late in infection, the symptoms worsen, and the infected individual develops dyspnea, pulmonary congestion, severe respiratory distress, hemoptysis, and ultimately shock [4, 5].

*Bacillus anthracis* is of particular interest to the biodefense sector since it is one of the few biological agents that has been released or used on civilians as a nefarious act of bioterrorism [6–8]. The World Health Organization has estimated that an anthrax spore release in a city of 5 million could cause up to 250,000 casualties with 100,000 deaths [9]. Given this threat, the US Department of Health and Human Services is tasked with stockpiling medical countermeasures and ensuring “timely and accurate recommended utilization guidelines” to protect the public against anthrax [10]. The ensuing mortality and morbidity would be further compounded with the use of *B. anthracis* strains that were genetically modified to be resistant to antibiotics approved by the Food and Drug Administration (FDA) currently used to treat anthrax infection, such as ciprofloxacin, doxycycline, and levofloxacin. Due to this potential threat, there remains a need for new antibiotic treatments to combat the disease. Such novel medical countermeasures (MCMs) should be developed and added to the Strategic National Stockpile as current MCM stocks demonstrate less or no activity as first-line antibiotic therapy in the event of a bioterrorist attack and resistant strains.

Telavancin (TD-6424, trade name Vibativ) is the only clinically approved semisynthetic glycopeptide antibiotic derived from vancomycin. It differs most significantly from its parent structure by the decylaminoethyl modification on the vancosamine unit, a modification that is responsible for telavancin’s enhanced potency against Gram-positive strains [11, 12]. This modification introduced unfavorable excretion and distribution properties, so an additional (phosphonomethyl) aminomethyl moiety was appended to ring 7, leading to an improved ADME profile [11, 12].

Telavancin has a dual mode of action. First, it retains the mechanism of action of vancomycin by binding lipid II, thereby inhibiting bacterial cell wall biosynthesis [13, 14]. This interaction is promoted by the decylaminoethyl lipid, which anchors into the cytoplasmic membrane and brings telavancin into proximity with peptidoglycan precursors. For this reason, telavancin displays a higher binding affinity for the bacterial cell surface and increased inhibition of transglycosylation [14]. Telavancin’s lipid moiety is responsible for the second mode of action: the concentration-dependent dissipation of bacterial cell membrane potential (at 10-fold MIC), leading to membrane permeabilization and leakage of ATP and potassium ions [13–15]. Telavancin displays a low propensity to induce spontaneous resistance in staphylococci and enterococci [16]. Interestingly, this dual mechanism of action gives telavancin a potency 10-fold greater than vancomycin [17–19].

In the US and Canada, telavancin was approved in September 2009 for use in the treatment of complicated skin and skin structure infections (CSSSIs) caused by susceptible Gram-positive species such as *S. aureus*, *Streptococcus agalactiae*, *Streptococcus pyogenes*, and *Enterococcus faecalis*, and in June 2013 for use in case of hospital-acquired pneumonia including ventilator-associated pneumonia caused by *S. aureus* [20]. Telavancin is active against a variety of Gram-positive species, including MRSA (MIC = 0.016–0.125 μg/mL), VanB-type VRE (MIC = 2 μg/mL), and *S. pneumoniae* (MIC = 0.008–0.03 μg/mL) [21]. Telavancin’s MICs against vancomycin-resistant *E. faecalis* and *E. faecium* are ≤0.025–6 mg/l and 0.015–16 mg/l, respectively. For 29 isolates of vancomycin-resistant *E. faecium* as well as 29 isolates of vancomycin-resistant *E. faecalis*, the MIC_90_ (MIC needed to inhibit 90% strain growth) is 1/64 (∼0.015625) times the MIC_90_ of vancomycin (viz., 4 mg/l *versus* >256 mg/l) [22]. Many species of streptococci (including multidrug-resistant and penicillin-resistant strains) and *Listeria* (MIC, 0.125 mg/l) are susceptible to telavancin [22–24]. Unlike teicoplanin, telavancin is also potent against VISA strains [17, 25]. Telavancin is more active against VISA and hVISA than vancomycin, given that the MIC_90_ of vancomycin is 8 times more than that of telavancin (1 mg/l *versus* 8 mg/l) for 50 isolates of glycopeptide-intermediate staphylococcus species (GISS) and heteroresistant GISS [22]. Telavancin is known to exert activity against *S. aureus* harbored intracellularly inside murine THP-1 and human J774 macrophage cell lines [26]. Biofilm-generating *S. aureus* and *Staphylococcus epidermidis* are also susceptible to the antibacterial effects of telavancin [27]. *Actinomyces* species (MIC, 0.125–0.25 mg/l), *Clostridium difficile* (MIC, 0.125–0.5 mg/l), and many other anaerobic bacteria are susceptible to telavancin [28].

The pharmacokinetic-pharmacodynamic profile of telavancin allows for once-daily dosing with adequate penetration into the skin and lungs and no requirement for monitoring serum drug concentrations. The manufacturing process has established a reliable telavancin supply for inpatient and outpatient parenteral antimicrobial therapy. Telavancin was tested against *B. anthracis* in two separate *in vitro* experiments. The first tested 15 strains [29], and the current study tested 17 strains. Both demonstrate that telavancin has potent activity against all *B. anthracis* strains with MIC at or below 0.125 ug/ml.

These results led us to test the protective efficacy of telavancin against inhalation anthrax infection using the rabbit model. The *in vivo* study was performed using the rabbit model, which is a superior model for inhalation anthrax infection relative to rodent models [30] and was conducted as a trigger-to-treat study; therefore, treatment began after *B. anthracis* protective antigen (PA) was detected in each animal’s sera via electrochemiluminescence. Blood and tissue samples were also collected to assess the extent of infection and the effectiveness of telavancin. To our knowledge, this is the first reported work showing the therapeutic potential of telavancin against inhalation anthrax in a large animal model.

## Results

### Telavancin *in vitro* activity against *B. anthracis* strains

Telavancin was screened against a diverse panel of *B. anthracis* strains consisting of laboratory and clinical strains considered representative of the drug susceptibility spectrum associated with clinical infections [31]. Telavancin has a minimal inhibitory concentration range of 0.0625-0.125 mg/L against the laboratory reference strain and the panel of clinical strains with various susceptibilities to standard-of-care drugs (**Table 1**). This MIC value and inhibition are characteristic of clinical drugs with bactericidal activity against *B. anthracis*, which is within the range for a drug to have efficacy. Most significantly, telavancin demonstrated superior potency to current clinically used drugs to treat *B. anthracis* infections with no observable spontaneous drug resistance. For example, the MICs determined for various clinical strains of *B. anthracis* with doxycycline and vancomycin fall into a range of 0.0156-0.0312 mg/L and 1-4 mg/L, respectively, which is comparable to other clinical drugs [31] and represents a significant difference in the selective index (SI) between tissue toxicity and bactericidal dose.

**Table 1:** MICs summary for telavancin and doxycycline against B. doxycycline against B. anthracis strains

| Strain | Telavancin | Doxycycline |
| --- | --- | --- |
| Ames 3838 | <0.0625 | 0.03125 |
| Graves | <0.0625 | 0.03125 |
| 46-PY-5 | <0.0625 | 0.03125 |
| Ames | <0.0625 | 0.03125 |
| Kruger B (A0442) | 0.125 | 0.03125 |
| Vollum (A0488) | <0.0625 | <0.0156 |
| WNA | <0.0625 | 0.03125 |
| A0318 | <0.0625 | 0.03125 |
| A0471 | <0.0625 | 0.03125 |
| Anthrax Strain Collection (ASC) 506 | <0.0625 | 0.03125 |
| Anthrax Strain Collection (ASC) 525 | <0.0625 | 0.03125 |
| 2000032823 (CDC #1) | <0.0625 | <0.0156 |
| 2002734753 (CDC #2) | 0.125 | 0.03125 |
| 2010719149 (CDC #3) | <0.0625 | 0.03125 |
| 2006200760 (CDC #4) | <0.0625 | <0.0156 |
| ASC 32 | <0.0625 | 0.03125 |
| ASC 149 | <0.0625 | 0.03125 |

### Infection and Toxemia

The mean aerosol exposure dose of *B. anthracis* spores for 30 animals was 2.25 x 10^7^ CFU (± 4.03 x 10^5^), corresponding to 225 LD_50_ (50% lethal dose), and the mean duration of the aerosol challenges was 13 min. Analysis of sera extracted from whole blood samples collected every 6 hrs beginning 12 hrs post-infection showed exposed animals exhibited toxemia (as measured by detectable levels of PA using an ECL assay) as early as 18 hrs post-infection and as late as 30 hrs post-infection (7%). In comparison, the majority (63%) of the animals were positive for PA at 18 hrs post-infection (**Table 2**). The concentration of PA in the sera ranged from approximately 50 to 2,000 picograms per milliliter (pg/ml).

**Table 2.** Time points of PA detection

| Treatment group | Animal ID# | Time of PA detection | PA concentration (pg/ml) |
| --- | --- | --- | --- |
| Telavancin | 8629 | 18 | 960 |
|  | 8630 | 18 | 597 |
|  | 8633 | 18 | 824 |
|  | 8637 | 18 | 371 |
|  | 8639 | 18 | 796 |
|  | 8641 | 24 | 348 |
|  | 8643 | 18 | 1,781 |
|  | 8646 | 24 | 2,039 |
|  | 8647 | 24 | 639 |
|  | 8651 | 18 | 418 |
|  | 8653 | 24 | 841 |
|  | 8656 | 18 | 377 |
| **Levofloxacin** | 8628 | 18 | 1,052 |
|  | 8632 | 18 | 1,128 |
|  | 8634 | 18 | 58.4 |
|  | 8635 | 24 | 468 |
|  | 8638 | 18 | 146 |
|  | 8640 | 18 | 154 |
|  | 8642 | 24 | 514 |
|  | 8644 | 30 | 258 |
|  | 8649 | 24 | 1,189 |
|  | 8652 | 18 | 1,229 |
|  | 8654 | 18 | 146 |
|  | 8657 | 24 | 269 |
| **Saline** | 8631 | 18 | 509 |
|  | 8636 | 30 | 1,007 |
|  | 8645 | 18 | 922 |
|  | 8648 | 18 | 389 |
|  | 8650 | 18 | 746 |
|  | 8655 | 24 | 164 |

### Survival

Following infection, the animals were monitored at least twice daily for 14 days. The most common clinical signs included anorexia, lethargy, and respiratory distress. These observations were consistent with symptoms of inhalational anthrax. The telavancin-treated group exhibited 100% survival after the challenge which was significantly (p˂0.001) greater than that of the saline-treated group, which showed no survival (**Fig 1**). As expected, the animals administered saline succumbed to infection 2 to 4 days post-infection. The group treated with the humanized levofloxacin dose (12.5 mg/kg) also showed 100% survival. These results indicate that telavancin was completely protective in this model of inhalation anthrax infection.

**Fig 1.**
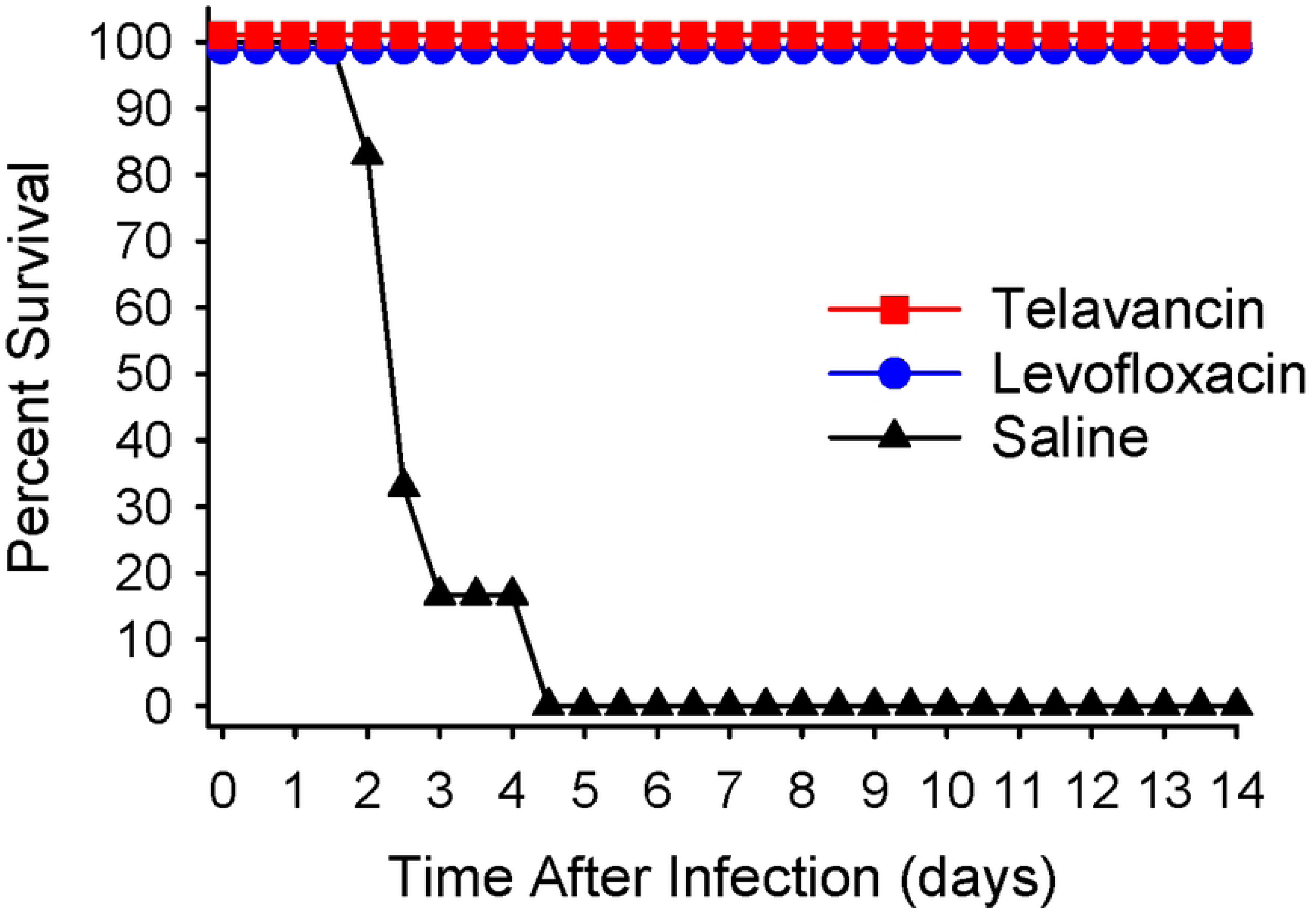
The therapeutic efficacy of telavancin and levofloxacin in the rabbit model of inhalation anthrax. New Zealand White rabbits were challenged with ∼200 LD_50_ *B. anthracis* Ames spores via inhalation. Telavancin treatment was initiated upon detection of PA in the animals’ sera and was administered at 30 mg/kg twice daily for 5 days. Levofloxacin, at 12.5 mg/kg, administered once daily for 5 days, and saline, administered once daily, were used as controls. Survival was monitored for 14 days post-infection. The percent survival rates for the two antibiotic-treated groups were significantly (p<0.001) higher than that of the saline-treated group.

### Temperature response during infection and treatment

The antibiotic-treated groups (telavancin and levofloxacin) exhibited comparable temperature responses after the challenge (**Fig 2**). Specifically, animals in both groups had febrile responses at approximately Day 1 post-infection that peaked at nearly 41⁰C before Day 2. By Day 2, the mean temperatures for the antibiotic-treated groups returned to baseline, most likely due to treatment which began at 18 to 30 hrs post-infection. Interestingly, both antibiotic-treated groups appeared to have minor secondary febrile responses from 7 to 9 days post-infection, which was after treatment was ended (Day 6), but these temperature elevations subsided by Day 10 to 11. The mean temperature of the animals treated with saline also began to rise by Day 1, and it remained elevated until hours before the animals succumbed to infection. These results show that post-exposure treatment with telavancin was effective at reducing and resolving the febrile response consistently reported with anthrax infection.

**Fig 2.**
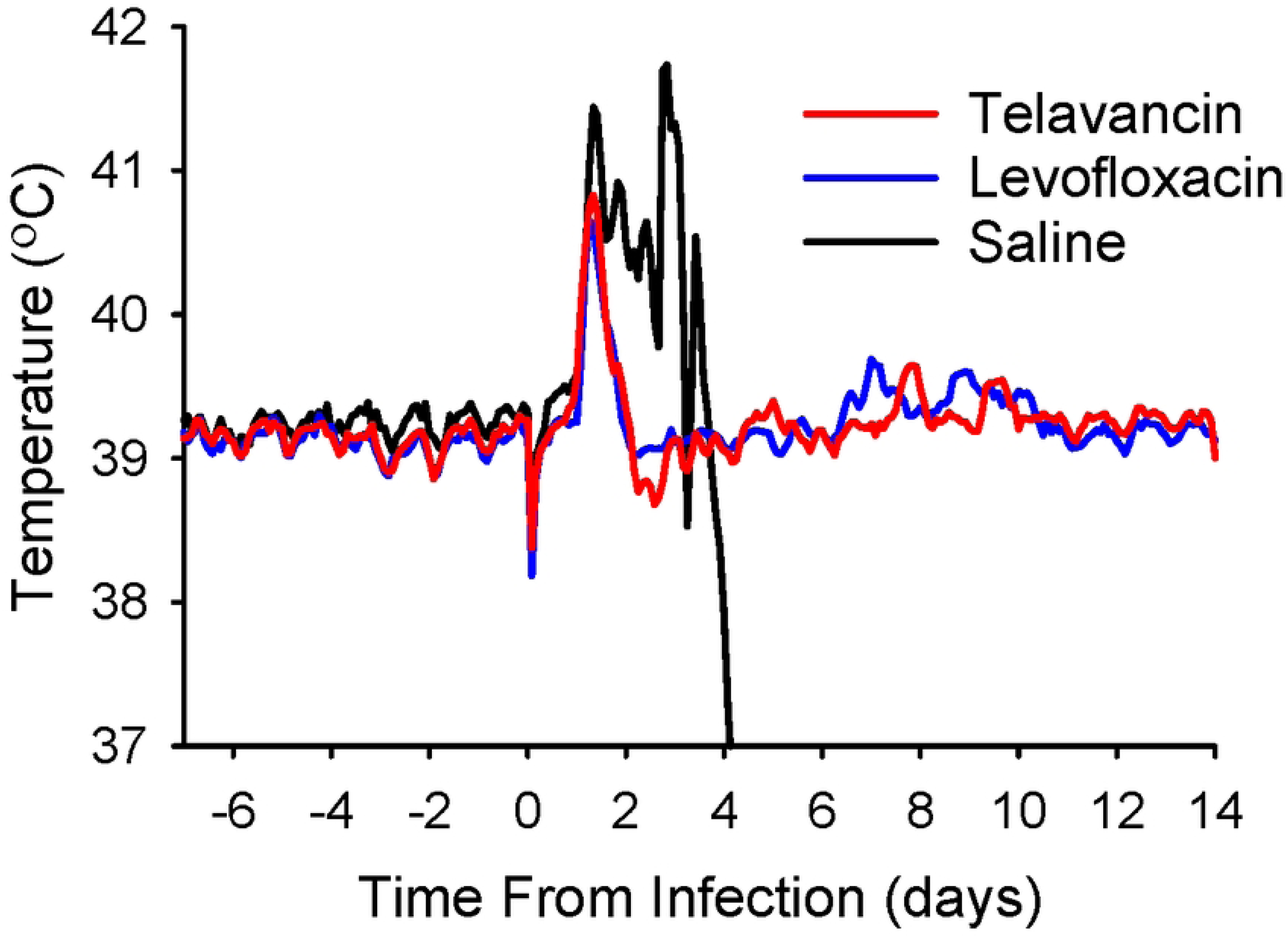
Temperature response during infection and antibiotic treatment. New Zealand White rabbits were challenged with 200 LD_50_ *B. anthracis* Ames spores via inhalation. Telavancin treatment was initiated upon detection of PA in the animals’ sera and was administered at 30 mg/kg twice daily for 5 days. Levofloxacin, at 12.5 mg/kg, administered once daily for 5 days, and saline, administered once daily, were used as controls. Temperatures were recorded every 10 min for 7 days before infection and for the entire post-infection period using implantable data loggers. The data are presented as two-hour moving averages.

### Bacteremia

**Table 3** indicates the *B. anthracis* bacterial load in the blood of each animal in the three treatment groups. The data are presented in colony forming unit per milliliter (cfu/ml). Bacteremia was detected as early as 24 hrs post-infection. At that time, the average levels of bacteremia for the antibiotic-treated (telavancin and levofloxacin) groups were significantly (p˂0.05) lower than that of the saline-treated group (**Fig 3**). Approximately half the animals in both the telavancin- and levofloxacin-treated groups received their initial treatment by 24 hrs post-infection. Interestingly, only 2 out of 12 animals treated with telavancin were positive at this time point but 8 out of 12 animals treated with levofloxacin were positive at this same time. Moreover, when comparing the two antibiotic-treated groups, the average level of bacteremia was significantly (p˂0.10) lower among the animals treated with telavancin (**Fig 3**). This would suggest that the humanized telavancin dosage was more effective at eliminating the bacteria in the blood relative to the humanized levofloxacin dosage used. By 48 hrs post-infection, all antibiotic-treated animals were negative for bacteremia and remained so for the remainder of the post-infection period (Days 7, 10, and 14 post-infection, not shown). Overall, these results demonstrate that rabbits treated with telavancin more rapidly clear *B. anthracis* from circulation than rabbits treated with levofloxacin.

**Fig 3.**
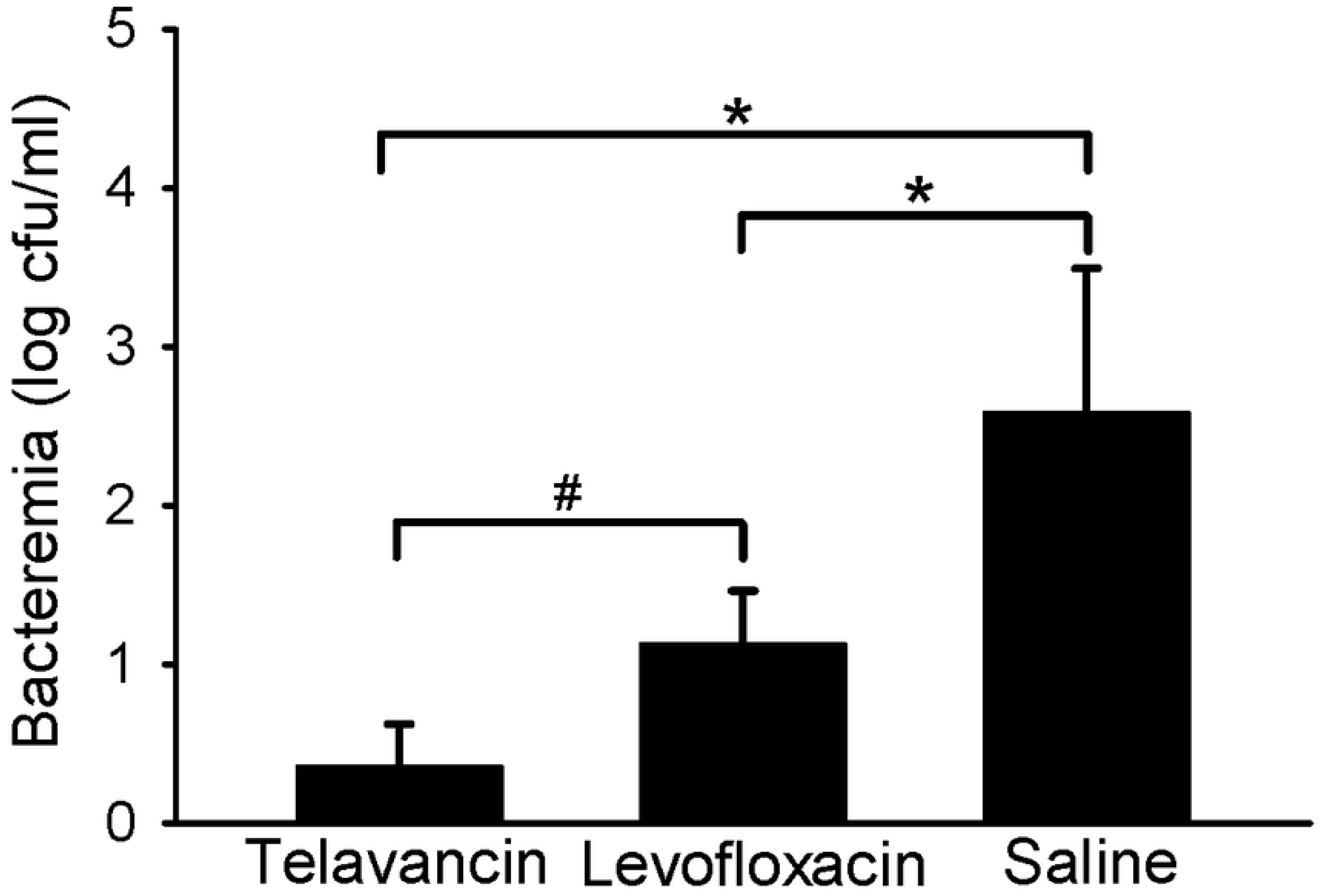
*B. anthracis* bacteremia levels during infection and antibiotic treatment at 24 hours post-infection. New Zealand White rabbits were challenged with 200 LD_50_ *B. anthracis* Ames spores via inhalation. Telavancin treatment was initiated upon detection of PA in the animals’ sera and was administered at 30 mg/kg twice daily for 5 days. Levofloxacin, at 12.5 mg/kg, was administered once daily for 5 days, and saline, administered once daily, was used as a control. Whole blood was plated on TSAII plates and incubated at 37°C for 16 to 24 hrs. Data are presented as averages with standard error bars. The asterisk and hashtag indicate significant (p<0.05 and p<0.10, respectively) differences between groups.

**Table 3.**
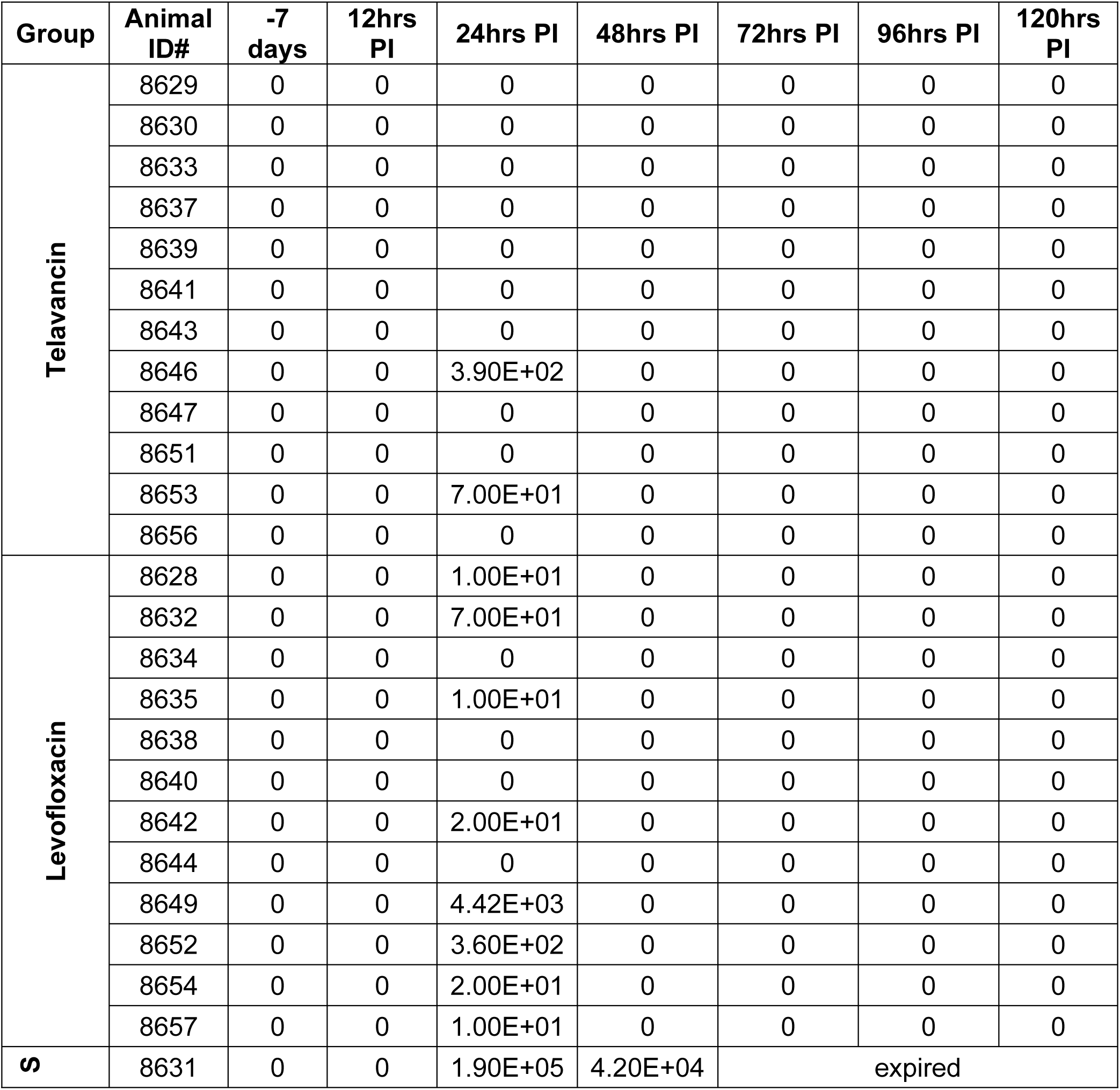
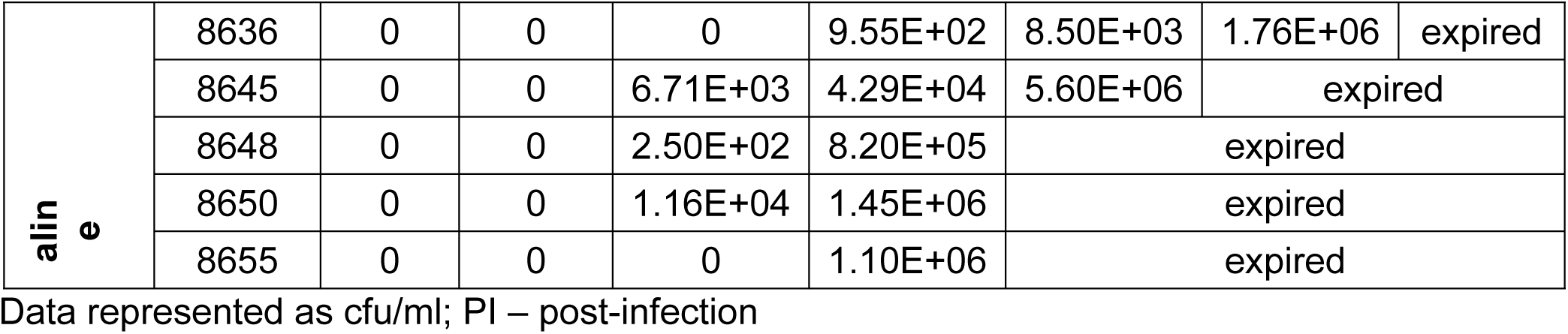
*B. anthracis* bacteremia levels

### Bacterial load in tissues

**Table 4** gives the *B. anthracis* bacterial load in lung tissue collected from each animal in the three treatment groups at the time of euthanasia due to reaching humane or scientific endpoints. Although *B. anthracis* was also detected in the brains, mediastinal lymph nodes and spleens of the saline-treated control animals (average bacterial loads of 6.68 x 10^5^ cfu/g, 2.59 x 10^6^ cfu/g, and 6.43 x 10^6^ cfu/g, respectively), no bacteria were recovered from these tissues from any animal in either group treated with telavancin or levofloxacin. In a comparison of *B. anthracis* bacterial loads in lung tissue, there were significantly (p<0.05) fewer bacteria in the two antibiotic-treated groups relative to the saline control group (**Fig 4**). When comparing the two antibiotic-treated groups, the average bacterial load in the lung tissue of the animals treated with telavancin (1.93 x 10^3^ cfu/g) was significantly (p˂0.05) lower than that of the animals treated with levofloxacin (6.93 x 10^3^ cfu/g) (**Fig 4**), suggesting once again that the humanized telavancin dosage was more effective at eliminating the bacteria in tissues relative to the humanized levofloxacin dosage. Since tissues were not collected for assessment of bacterial load at various scheduled times post-infection, it is unknown whether telavancin completely prevented the migration of *B. anthracis* to the tissues or if it cleared the bacteria after the bacteria entered the tissues. Nonetheless, these results show that telavancin was highly efficacious compared to levofloxacin in preventing, clearing, and reducing *B. anthracis* infection of tissues after exposure to inhalation anthrax.

**Fig 4.**
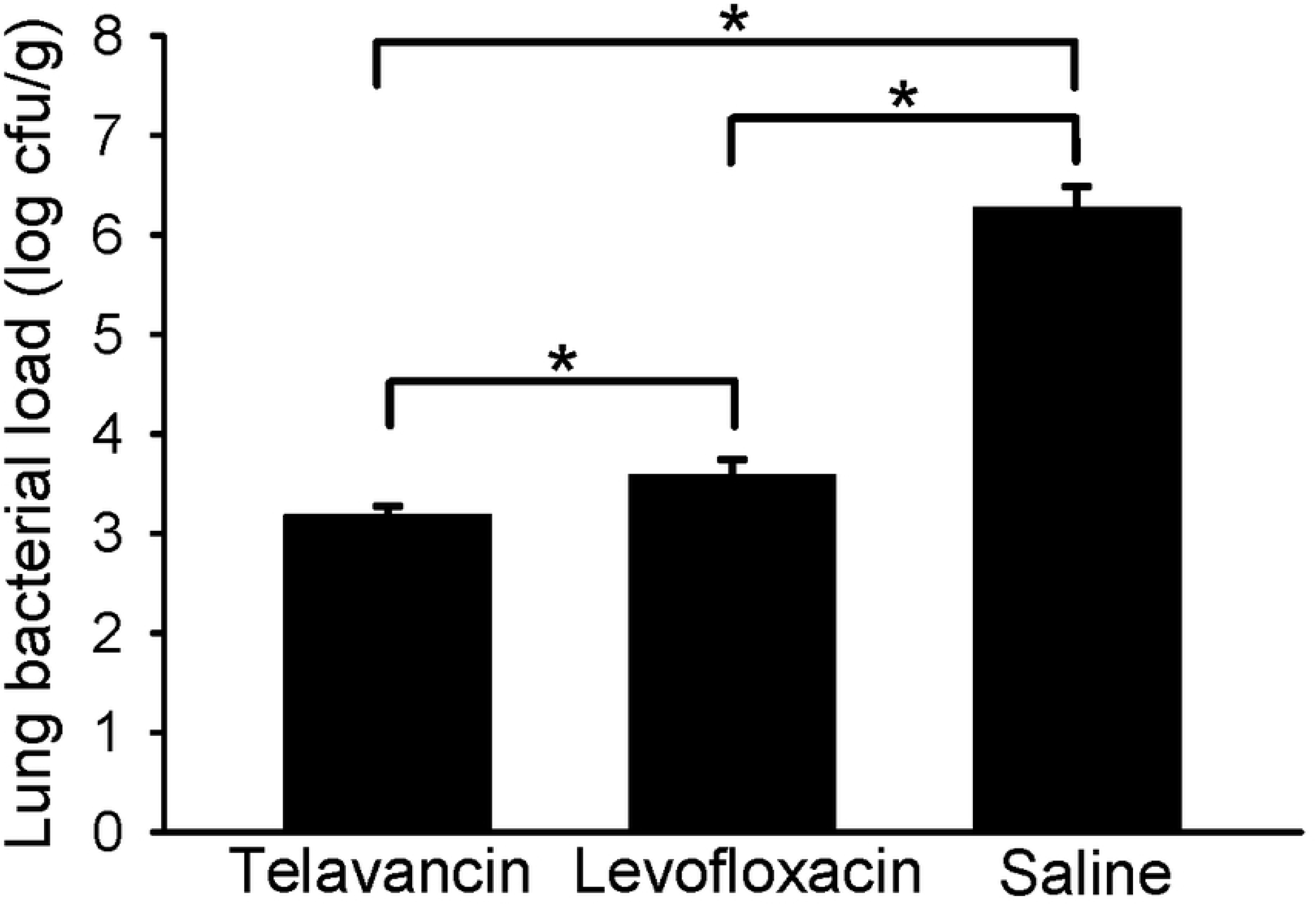
*B. anthracis* bacterial load in lung tissue at time of euthanasia. New Zealand White rabbits were challenged with 200 LD_50_ *B. anthracis* Ames spores via the inhalation route. Telavancin treatment was initiated upon detection of PA in the animals’ sera and was administered at 30 mg/kg twice daily for 5 days. Levofloxacin, at 12.5 mg/kg, and 0.9% sodium chloride solution, were each administered once daily for 5 days and served as the positive and negative controls, respectively. Tissues were homogenized, and the homogenates were plated on TSAII plates. The plates were then incubated at 37°C for 16 to 24 hrs. Data are presented as averages with standard error bars. The asterisk indicates a significant (p<0.05) difference between groups.

**Table 4.**
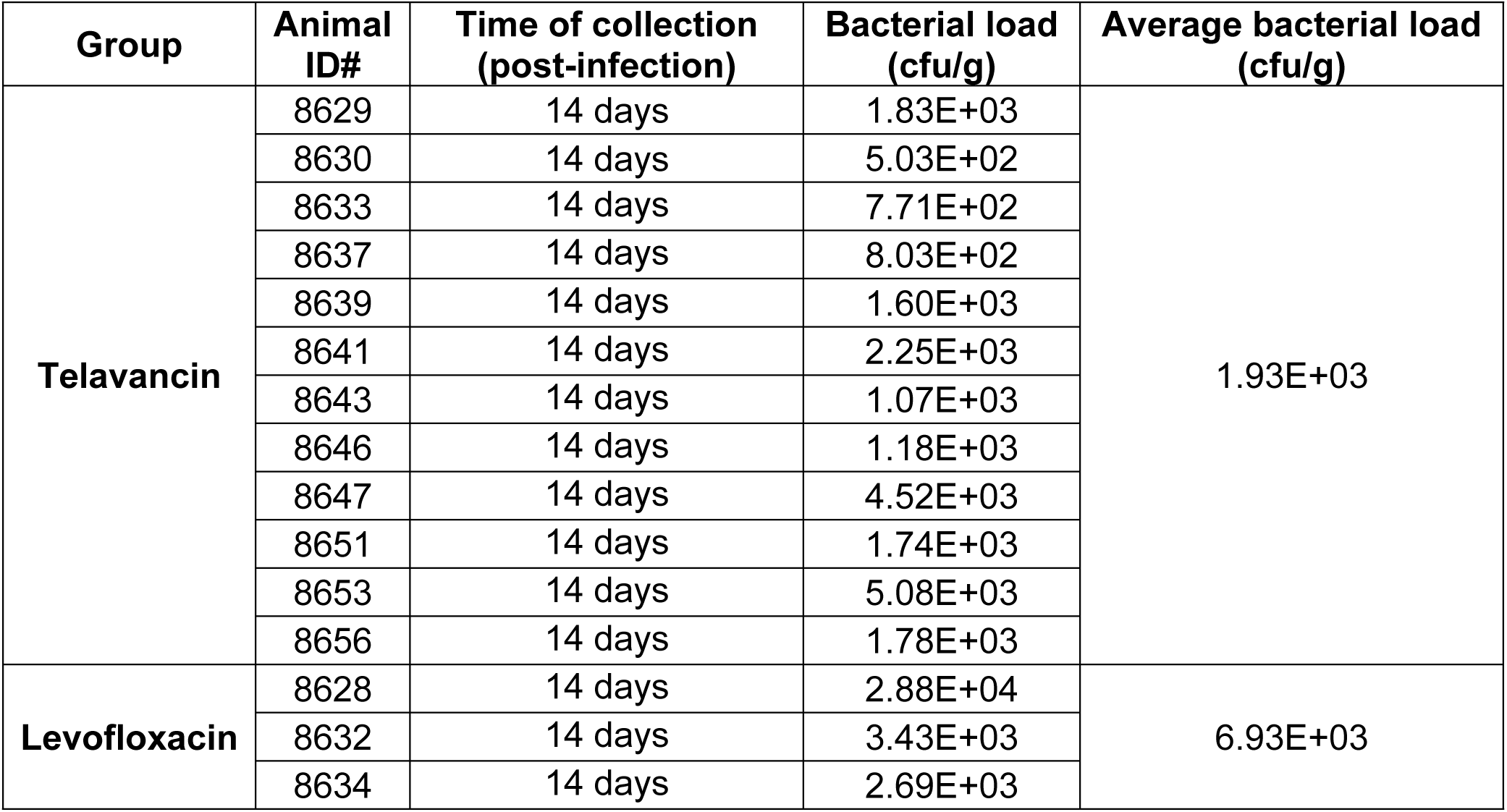
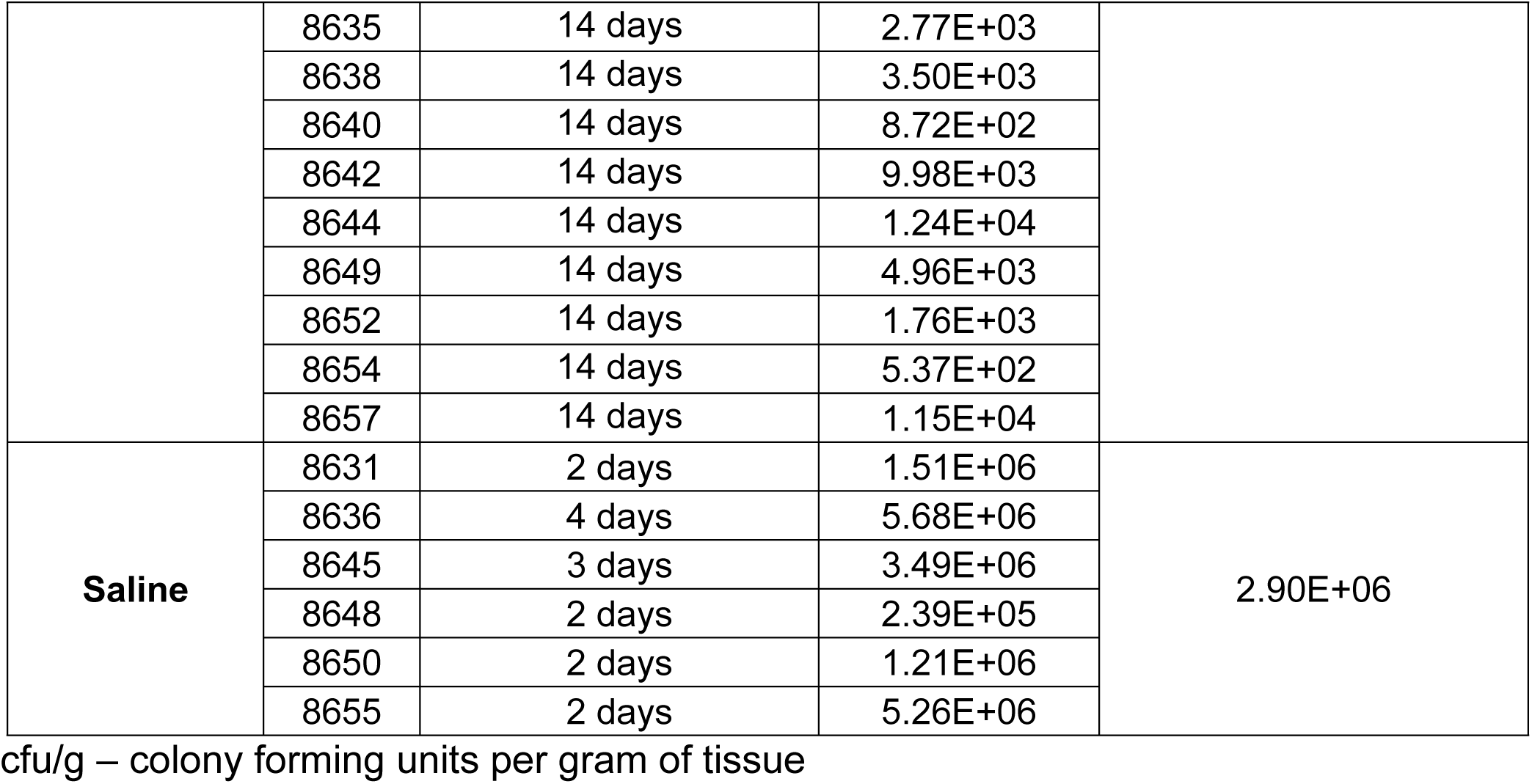
*B. anthracis* bacterial load in lung tissue

### Assessment of anti-PA antibody levels

The animals in the two antibiotic-treated groups began to develop antibody responses to *B. anthracis* protective antigen as early as 7 days post-infection, and the responses intensified at Days 10 and 14 post-infection (**Fig 5**). There was no significant difference in average serum anti-PA IgG titers between the telavancin- and levofloxacin-treated groups; however, both antibiotic-treated groups had a significant increase (p<0.05) over time. On the other hand, the negative control group showed no increase in anti-PA IgG levels over time since they succumbed to the anthrax infection before the immune response could produce significant levels of serum anti-PA IgG. These results demonstrate that telavancin did not alter the humoral immune response to PA generated following antibiotic treatment of anthrax infection.

**Fig 5.**
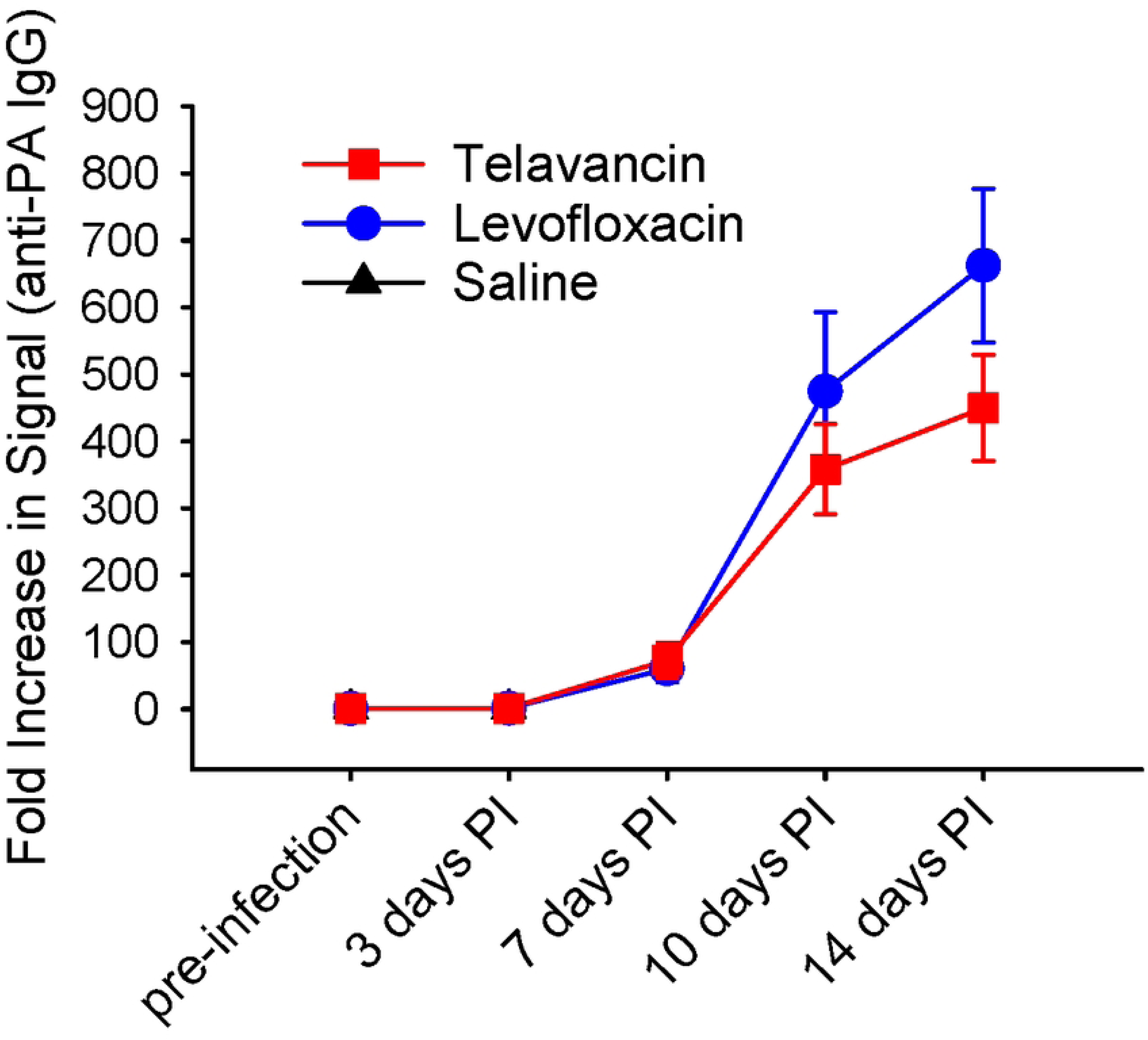
Anti-PA antibody response during infection and antibiotic treatment. New Zealand White rabbits were challenged with 200 LD_50_ *B. anthracis* Ames spores via inhalation. Telavancin treatment was initiated upon detection of PA in the animals’ sera and was administered at 30 mg/kg twice daily for 5 days. Levofloxacin, at 12.5 mg/kg, was administered once daily for 5 days, and saline, administered once daily, was used as controls. Sera were extracted from whole blood at various time points, and the anti-PA antibody response was measured in the sera via ECL. Data are presented as averages with standard error bars; PI = post-infection.

### Necropsy and Histopathology

At necropsy, the animals in the saline-treated group consistently showed blackened, enlarged, hemorrhagic mediastinal lymph nodes, discoloration, and diffuse hemorrhage of the lungs with pleural effusion. There was also pericardial effusion and multifocal hemorrhage on the serosal surface of the vermiform appendix. Additional notable findings, although less consistent, in the saline-treated group were the presence of hemorrhage in the meninges, multifocal discoloration of the cecum, and hemorrhage in the thymus. Importantly, all these findings have been reported to be associated with anthrax infection in animal models. The gross pathology observed among the groups treated with telavancin, and levofloxacin was far less prominent than in the untreated control animals. Animals in these antibiotic-treated groups still exhibited a few pathologic findings, such as pulmonary congestion, multifocal hemorrhage of the lungs, and enlarged mediastinal lymph nodes, indicating an active infection, but the lesser pathology among these groups relative to the untreated controls serves as evidence of disease mitigation.

Histopathological analysis showed notable lesions in the tissues of the control animals, with the lesions in the lungs, mediastinal lymph nodes, and spleen being the most significant (**Fig 6**). Edema was consistently present in the lungs/alveoli of the control animals, and bacteria were seen intermittently in the alveoli vasculature. The mediastinal lymph nodes showed diffuse degeneration and necrosis along with regions of hemorrhage. Similarly, the spleen showed diffuse necrosis (**Fig 6**). The remaining collected tissues in the control animals also exhibited pathology, albeit to a lesser extent. Specifically, there was evidence of degeneration and necrosis in the heart and liver; bacteria were present in the liver, kidneys, and brain, and hemorrhage was present in the brain (**S1 Fig**). Conversely, tissues from animals treated with telavancin and levofloxacin showed only evidence of disease mitigation, and there were no substantial differences among the tissues from these two groups (**Fig 6** **and S1 Fig**). These results show that telavancin treatment effectively clears *B. anthracis* from tissues, ultimately leading to disease mitigation and survival.

**Figure 6.**
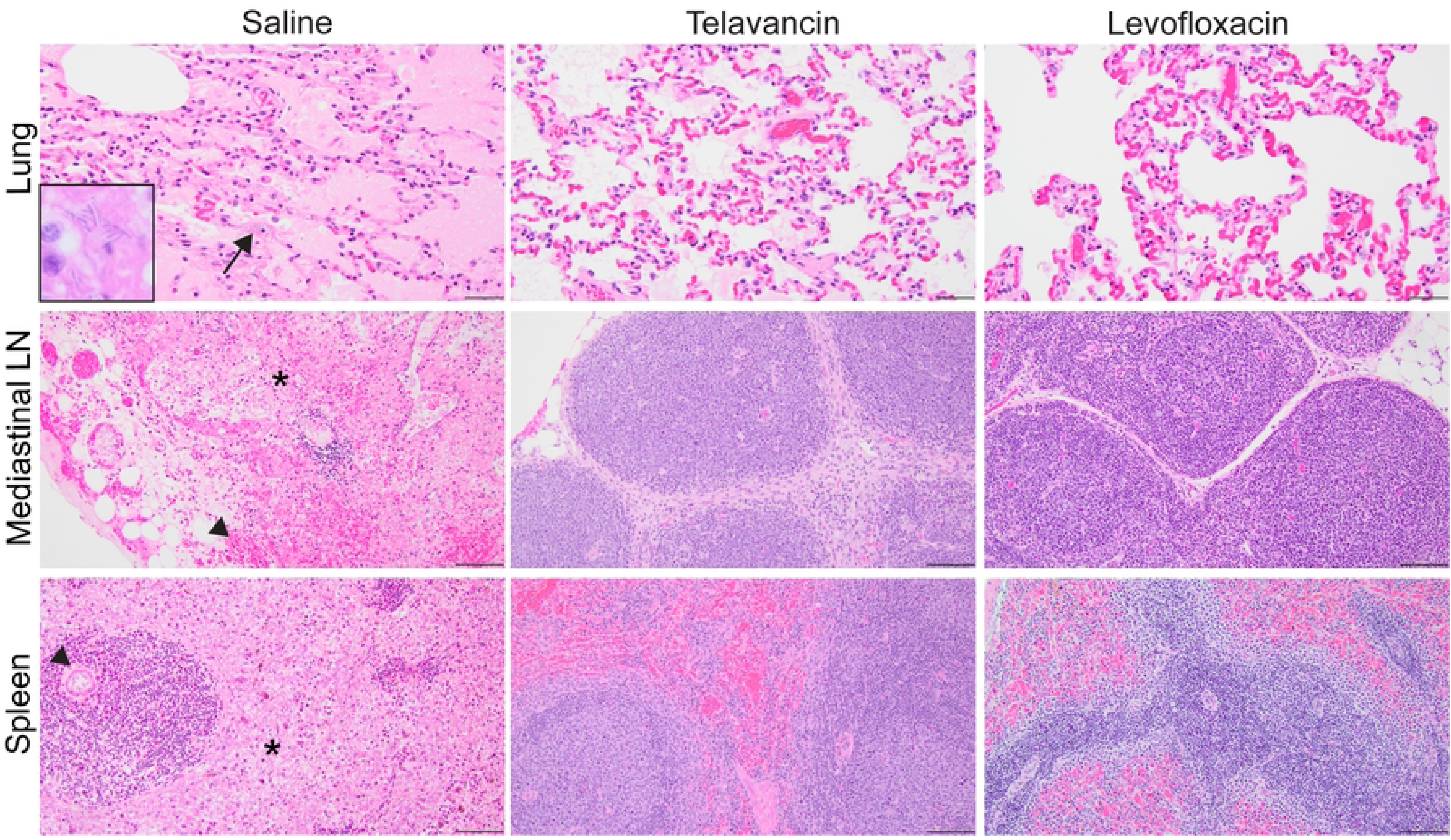
Histopathological lesions in lungs, spleen, and lymph nodes of rabbits infected with inhalation anthrax and treated with telavancin and levofloxacin. New Zealand White rabbits were inoculated with anthrax via the inhalation route. Lung, mediastinal lymph node, and spleen were collected for routine histopathology. Significant lesions were only detected in the animals dosed with saline. Lung: The alveoli contain edema fluid. Bacteria are within the alveoli vasculature (arrow). Inset, higher magnification of intravascular anthrax bacilli. Mediastinal lymph node: Diffusely, the lymphoid follicles are necrotic (*) with areas of hemorrhage (arrowhead). Spleen: The white pulp (arrowhead), and red pulp (black asterisk) are diffusely necrotic. Hematoxylin and eosin. Scale bar = 20 µm.

## Discussion

Telavancin (TD-6424) is a lipoglycopeptide derivative of vancomycin administered once daily as a 1-hour infusion at a dose of 10 mg/kg/day for the treatment of CSSSIs and nosocomial pneumonia (hospital-acquired and ventilator-associated) in adults with a CrCl >50 mL/min. Telavancin has excellent bactericidal activity against aerobic and anaerobic Gram-positive bacteria, including multiple resistant phenotypes of *S. aureus,* and superior efficacy compared to vancomycin in diverse animal models of difficult-to-treat gram-positive infections, such as pneumonia, bacteremia, and endocarditis [17,20,32–34]. Although the bactericidal activity of telavancin against many clinically relevant Gram-positive bacteria has been demonstrated [25, 35], limited data are available on *B anthracis*. Here, we investigated the *in vitro* activities of telavancin against 15 strains of *B. anthracis*. In our study, telavancin exhibited potent bactericidal activity against the 15 *B anthracis* strains tested with MICs <0.0625 μg/mL for all but 2 strains with MIC of 0.125 (Kruger B [A0442] and 2002734753 [CDC #2]) similar to the FDA-approved and comparator agent, doxycycline and levofloxacin, parenteral procaine penicillin G and ciprofloxacin. Notably, the susceptibility profile of these diverse strains demonstrated resistance and intermediate susceptibility across multiple families of antibiotics [36] and combinations, underscoring the importance of expanding the spectrum of antibiotics available for the treatment of anthrax.

These encouraging *in vitro* results led us to assess the protective efficacy of telavancin *in vivo*. In the present study, telavancin was shown to be completely protected against inhalation anthrax infection using New Zealand White rabbits challenged with approximately 200 LD_50_ *B. anthracis* Ames spores. Moreover, this study was conducted as a trigger-to-treat study wherein telavancin treatment was not initiated until the infection was confirmed by detection of the *B. anthracis* PA. This aspect is beneficial for real-time application since anthrax infection presents nonspecific symptoms that make clinical diagnosis difficult.

Results showed that telavancin could quickly eliminate *B. anthracis* from circulation, as evidenced by the absence of bacteremia and the waning fever response one day after start of treatment. Interestingly, the level of bacteremia at 24 hrs post-infection was lower for the telavancin-treated group compared to our positive control, the levofloxacin-treated group, which would suggest that telavancin was more effective than levofloxacin in eliminating the bacteria; Telavancin was also able to clear the bacteria more effectively from select tissues compared to levofloxacin, although, tissue clearance is partly due to an animal’s own acquired immunity since tissues of the surviving animals were harvested 14 days post-infection. A robust anti-PA IgG antibody response was evident 7 to 10 days post-infection. *B. anthracis* was, for instance, detected in the lungs of both antibiotic-treated animals at termination. However, the bacterial load was significantly less than that of the saline-treated animals (**Fig. 4**). Interestingly, the telavancin-treated group had a lower bacterial load in the lungs compared to the levofloxacin group, which would suggest once again that telavancin was more effective than levofloxacin at eliminating the bacteria. The anthrax spores persisted in the lungs for the entire post-infection period, but the prolonged presence of spores in the lungs after an initial exposure is a characteristic feature of inhalation anthrax. In fact, dormant anthrax spores have been recovered from the lungs of nonhuman primates [37] and mice [38, 39] weeks or months post-exposure. This is a possible explanation for this study’s mild secondary fever responses observed among the antibiotic-treated groups. Based on these observations, the CDC recommends administering post-exposure prophylaxis antimicrobial therapy for 60 days [40].

The CDC recently sponsored and published an Anthrax Preparedness supplement [41] with over a dozen articles in response to bioterrorism concerns, given the potential for *B. anthracis* to be weaponized. Since prospective studies in humans are limited, the CDC evaluated the effectiveness of various classes and combinations of antimicrobials by reviewing the outcomes of over 600 patients hospitalized with anthrax published over the last century [42] and concluded combination therapy to be superior to monotherapy for inhalation anthrax; however, neither monotherapy nor combination treatment is effective against anthrax meningitis. These findings highlight the unmet need for anthrax meningitis and *in vivo* studies to evaluate new antimicrobials and determine whether these new and current anti-anthrax agents penetrate the blood-brain barrier to be effective treatments for anthrax meningitis. The efficacy of telavancin (30 mg/kg) was evaluated in a rabbit model of meningitis caused by a strain of penicillin-resistant pneumococcus and compared with a combination of ceftriaxone (100 mg/kg) and vancomycin (20 mg/kg). Telavancin produced a more rapid lowering of cerebrospinal fluid (CSF) titers (−0.84 CFU/ml.h) than co-administration of vancomycin and ceftriaxone (−0.61 CFU/ml.h) [43], and the CSF was sterilized in six out of ten rabbits by telavancin and only four out of ten rabbits in the comparator arm. Of note, it was reported that the bacteria load at the initiation of therapy was significantly higher in the telavancin arm. These results warrant further studies with telavancin in a similar model of anthrax meningitis. Currently, treatment studies for anthrax meningitis in animal models are lacking. Future anthrax meningitis studies with telavancin in primates would be the most applicable to humans.

The CDC’s retrospective outcome study also underscores efforts to evaluate new combination therapies for synergy will be critical for successful anthrax preparedness. The combination of telavancin administered along with piperacillin-tazobactam, cefepime, imipenem, or ciprofloxacin for infections caused by single isolates of MRSA, VISA, VRSA, *S. agalactiae*, vancomycin-susceptible *E. faecalis*, vancomycin-resistant *E. faecium*, and daptomycin-resistant *S. aureus* reported no antagonism. When combinations of telavancin + piperacillin-tazobactam or telavancin + imipenem were employed against isolates of VISA, synergistic activity was reported. A synergistic effect was also recorded when telavancin was administered along with piperacillin-tazobactam, cefepime, or imipenem against a VRSA isolate [44]. These findings strongly indicate a need to progress well-designed animal-model treatment studies with telavancin to determine which combination with other anti-anthrax agents result in improved survival.

In summary, telavancin has potent *in vitro* activity against *B. anthracis* and is protective against lethal inhalation anthrax infection in the rabbit model. This widely used model mimics the human sequela associated with anthrax infection. Moreover, telavancin rapidly prevented disease progression and dissemination of the bacteria to the bloodstream and tissues. With its long half-life in humans, dual mechanism of action which differs from currently approved anti-anthrax agents, and synergistic activity, telavancin would be an ideal candidate for use as a medical countermeasure to inhalation anthrax infection. The results obtained with telavancin support further study to determine the protective dose in the nonhuman primate model of *B. anthracis* infection.

## Materials and Methods

### B. anthracis strains

The 17 *B. anthracis* strains used in this study are those specified in the defined panels of Category A pathogens used for drug candidate evaluation in the NIAID drug screening [45] and included the Ames reference strain (NR3838) and clinically derived strains [31].

### Telavancin *in vitro* activity to *B. anthracis* strains

MICs were determined in triplicate by the broth microdilution method in cation-adjusted Mueller-Hinton broth (CAMHB) according to the methodology of the Clinical and Laboratory Standards Institute (CLSI) [46, 47]. Vibativ was dissolved in a formulation supplied by Cumberland Pharmaceuticals (Nashville, TN) at a concentration of 6.4mg/ml and diluted to 256 µg/ml in the first column of a 96-well plate (final concentration was 128µg/ml when diluted 1:2 with culture). After 16 to 20 h incubation at 37°C, the MICs were determined visually. The quality control strain *Escherichia coli* ATCC 25922 was tested in parallel for 16-18 hours.

### Animals

New Zealand White rabbits (Envigo, Denver, PA), half males and half females, weighing approximately 3.0-3.5 kg, were surgically implanted with venous access ports (VAP) by the vendor to facilitate the collection of blood samples and the intravenous administration of the treatments. Upon receipt, the animals were surgically implanted with DST micro-T temperature data loggers (Star-Oddi Ltd, Gardabaer, Iceland) to record the animals’ temperature during the study. The animals were housed in ventilated cages with ad libitum access to food and water, and the room was maintained on a 12-hour light-dark cycle.

### Ethics statement

All animal procedures were conducted in accordance with an animal use protocol approved by the University of Texas Medical Branch Institutional Animal Care and Use Committee (IACUC).

### Bacterial spores

*B. anthracis* Ames (NR3838) spores were grown in modified Schaeffer’s medium using a New Brunswick fermenter. After inoculation, the fermenter was operated with aeration, and the pH was maintained at pH 7.0-7.5 for approximately 4 days, after which the crude spores were harvested aseptically by centrifugation. The spores were then washed with sterile molecular-grade water. The spores were purified by density gradient centrifugation using sterile MD-76. Visual observation of the spores at 400x by phase-contrast microscopy during each step of purification was performed to ensure the production of a homogeneous suspension of highly refractile spores.

### Bacterial infection

New Zealand White rabbits were challenged by aerosol with approximately 200 LD_50_ (50% lethal dose) of *B. anthracis* Ames spores (2.0 x 10^7^CFU (colony forming units)) using a Biaera aerosol control platform (Biaera Technologies, Hagerstown, MD) fitted with a head-only aerosol exposure chamber. Real-time plethysmography (Data Sciences International, St. Paul, MN) was performed on each animal during the aerosol exposure to monitor respiration. A 6-jet Collison nebulizer (CH Technologies, Westwood, NJ) was used to generate the aerosol. Aerosol samples were collected during each aerosol run using aerosol BioSamplers (SKC, Eighty-Four, PA) to confirm the challenge dose of spores for each animal by serial dilution and plating onto trypticase soy II agar plates containing 5% sterile sheep blood (TSAII). The duration of the aerosol delivery was based on the animal’s respiration and the total volume of inspired air.

### Antibiotic treatment

Telavancin (Cumberland Pharmaceuticals, Nashville, TN), provided as a lyophilized powder, was reconstituted in sterile, pyrogen-free water to a concentration of 25 mg/ml and administered intravenously to the animals at a dose concentration of 30 mg/kg twice daily for 5 days which mimics the levels measured in the serum of humans [48]. Telavancin treatments were prepared fresh daily. Levofloxacin (Akorn Pharmaceuticals, Lake Forest, IL) at a concentration of 25 mg/ml was administered intravenously to the animals at a clinically equivalent dose of 12.5 mg/kg once daily for 5 days. The untreated control animals were given saline once daily for 5 days. Treatments were initiated upon detection of protective antigen (PA) in the animals’ sera (point of antigenemia).

### Detection of PA

Using a rapid PA-electochemiluminescence (ECL) screening assay (MesoScale Discovery, Gaithersburg MD), the presence of PA was monitored in the animals’ sera that was extracted from whole blood (via the VAP) collected at various time points before and after infection. The time when PA was first detected was considered the point of antigenemia (PA in the blood) that dictated treatment initiation. A standard curve (0-100 ng/ml) was analyzed in parallel for each assay toto extrapolate the PA concentration of the sera samples. Test samples were assayed in duplicate.

### Assessment of bacteremia and bacterial load

The concentration of *B. anthracis* was measured in whole blood samples collected via the VAP at various time points before and after infection. Whole blood was plated onto TSAII plates using an automatic serial diluter and plater (Interscience Laboratories, Woburn, MA). After the plates were incubated at 37°C for 16-24 hrs, bacterial colonies were enumerated using an automatic colony counter (Interscience Laboratories, Woburn, MA). Bacterial colonies having morphology typical of *B. anthracis* were subcultured and confirmed as *B. anthracis* with bacteriophage ɣ.

Bacterial/spore load was determined in each animal’s lung, lymph node (mediastinal), brain, and spleen. These tissues were homogenized in sterile water using a Stomacher 80 MicroBiomaster (Seward Ltd, Bohemia, NY). The homogenates were serially diluted in water and plated onto TSAII plates using an automatic diluter/plater and incubated at 37°C for 16-24 hrs. Bacterial colonies were enumerated using an automated colony counter (Interscience Laboratories, Woburn, MA). Bacterial colonies having morphology typical of *B. anthracis* were subcultured and confirmed as *B. anthracis* with bacteriophage ɣ.

### Assessment of anti-PA antibody response

Anti-PA IgG was measured in serum via ECL, similar to the PA-ECL screening assay. Biotinylated recombinant PA83 (List Biological Labs, Campbell, CA) was bound to streptavidin-coated plates (MesoScale Discovery, Gaithersburg, MD) and used as the capture antigen. Detection was accomplished using SULFO-TAG labeled anti-rabbit antibody and read buffer (MesoScale Discovery, Gaithersburg, MD). Results were presented as a fold-increase in signal relative to the pre-challenge serum samples.

### Necropsy and histopathology

Gross pathology was performed on all animals, and tissues (lung, mediastinal lymph nodes, brain, and spleen) were collected and perfused with 10% phosphate-buffered formalin. Tissue sections were processed, embedded in paraffin, sectioned, stained with hematoxylin and eosin (H&E), and evaluated by microscopy. For analysis of the stained tissue sections, a four-level severity scale was used when applicable utilizing the following terms: minimal (1 of 4), mild (2 of 4), moderate (3 of 4), and marked (4 of 4).

### Statistical analyses

Statistical analyses were performed using NCSS (NCSS, Kaysville, UT), and differences between the experimental groups were tested using one-way Analysis of Variance (ANOVA) and Tukey-Kramer’s test.

## Acknowledgments

The authors would like to thank the veterinary staff of the University of Texas Medical Branch Animal Resources Center for their support in this project. We would also like to thank Breanne Gibson, Ph.D., for editing the manuscript.

## Supporting information

**S1 Fig. Histopathological lesions in liver, heart, kidneys, and brain of rabbits infected with inhalation anthrax and treated with telavancin and levofloxacin.** New Zealand White rabbits were inoculated with anthrax via the inhalation route. The liver, heart, kidney, and brain were collected for routine histopathology. Significant lesions were only detected in the animals dosed with saline. Liver: The liver has random areas of hepatocellular necrosis (asterisk) with intrasinusoidal bacteria (arrow). Inset, higher magnification of anthrax bacilli within the sinusoid. Heart: The cardiomyocytes are focally necrotic (asterisk). Kidney: The glomerular capillaries contain intravascular bacteria (arrow). Inset, higher magnification of anthrax bacilli within the capillary loops. Brain. Within the gray matter of the cortex are areas of hemorrhage (*) and intravascular bacteria (arrow). Inset, higher magnification of anthrax bacilli. Hematoxylin and eosin. Scale bar = 20 µm.

## Notes

### Competing Interest Statement

Ines Macias-Perez is an employee of Cumberland Pharmaceuticals.

## References

1. Beeching NJ, Dance DAB, Miller ARO, Spencer RC. Biological warfare and bioterrorism. BMJ. 2002;324:336–9.

2. U.S. Food and Drug Administration. Anthrax [Internet]. Anthrax. 2018 [cited 2023 Jan 25]. Available from: https://www.fda.gov/vaccines-blood-biologics/vaccines/anthrax#:~:text=The%20mortality%20rates%20from%20anthrax%20vary%2C%20depending%20on,a%20fatality%20rate%20that%20is%2080%25%20or%20higher.

3. Holty JEC, Brevata DM, Liu H, Olshen RA, McDonald KM, Owens DK. Systematic review: a century of inhalational anthrax cases from 1900 to 2005. Ann Intern Med. 2006 Feb;144(4):270–80.

4. Savransky V, Ionin B, Reece J. Current status and trends in prophylaxis and management of anthrax disease. Vol. 9, Pathogens. MDPI AG; 2020.

5. Kote CK, Welkos SL, Bozue J. Key aspects of the molecular and cellular basis of inhalational anthrax. Microbes Infect. 2011;13(14):1146–55.

6. Meselson M, Guillemin J, Hugh-Jones M, Langmuir A, Popova I, Shelokov A, et al. The Sverdlovsk Anthrax Outbreak of 1979. Science (1979). 1994;266:1202–7.

7. Eitzen E, Takafuji E. Historical overview of biological warfare. In: Sidell F, Takafuji E, Franz D, editors. Medical Aspects of Chemical and Biological Warfare. Washington, DC: Office of The Surgeon General at TMM Publications; 1997. p. 415–23.

8. Jernigan DB, Raghunathan PL, Bell BP, Brechner R, Bresnitz EA, Butler JC, et al. Investigation of Bioterrorism-Related Anthrax, United States, 2001: Epidemiologic Findings and the National Anthrax Epidemiologic Investigation Team 1. Vol. 8, Emerging Infectious Diseases. 2002.

9. World Health Organization Geneva. Health and Aspects of Chemical and Biological Weapons [Internet]. 1969 [cited 2023 Jan 25]. Available from: https://apps.who.int/iris/handle/10665/39444

10. 114th Congress. H.R.34 - 21st Century Cures Act [Internet]. 114th Congress; Dec 13, 2016. Available from: https://www.congress.gov/bill/114th-congress/house-bill/34

11. Judice JK, Pace JL. Semi-synthetic glycopeptide antibacterials. Bioorg Med Chem Lett. 2003 Dec 1;13(23):4165–8.

12. Leadbetter MR, Adams SM, Bazzini Kevin Krause BM, T Lam BM, Linsell MS, Kelly Quast M, et al. Hydrophobic Vancomycin Derivatives with Improved ADME Properties: Discovery of Telavancin (TD-6424). Vol. 57, JOURNAL OF ANTIBIOTICS. 2004.

13. Lunde CS, Hartouni SR, Janc JW, Mammen M, Humphrey PP, Benton BM. Telavancin disrupts the functional integrity of the bacterial membrane through targeted interaction with the cell wall precursor lipid II. Antimicrob Agents Chemother. 2009 Aug;53(8):3375–83.

14. Higgins DL, Chang R, Debabov D v., Leung J, Wu T, Krause KM, et al. Telavancin, a multifunctional lipoglycopeptide, disrupts both cell wall synthesis and cell membrane integrity in methicillin-resistant Staphylococcus aureus. Antimicrob Agents Chemother. 2005 Mar;49(3):1127–34.

15. Zhanel GG, Calic D, Schweizer F, Zelenitsky S, Adam H, Lagacé-Wiens PRS, et al. New Lipoglycopeptides A Comparative Review of Dalbavancin, Oritavancin and Telavancin. Drugs. 2010;70(7):859–86.

16. Kosowska-Shick K, Clark C, Pankuch GA, McGhee P, Dewasse B, Beachel L, et al. Activity of telavancin against staphylococci and enterococci determined by MIC and resistance selection studies. Antimicrob Agents Chemother. 2009 Oct;53(10):4217–24.

17. Draghi DC, Benton BM, Krause KM, Thornsberry C, Pillar C, Sahm DF. Comparative surveillance study of telavancin activity against recently collected gram-positive clinical isolates from across the United States. Antimicrob Agents Chemother. 2008 Jul;52(7):2383–8.

18. Hill CM, Krause KM, Lewis SR, Blais J, Benton BM, Mammen M, et al. Specificity of induction of the vanA and vanB operons in vancomycin-resistant enterococci by telavancin. Antimicrob Agents Chemother. 2010;54(7):2814–8.

19. Karlowsky JA, Nichol K, Zhanel GG. Telavancin: Mechanisms of Action, *in Vitro* Activity, and Mechanisms of Resistance. Clinical Infectious Diseases. 2015 Sep 15;61:S58–68.

20. Vibativ (telavancin) [package Insert]. Nashville, TN: Cumberland Pharmaceuticals, Inc; 2020.

21. European Committee on Antimicrobial Susceptibility Testing. The European Committee on Antimicrobial Susceptibility Testing. Breakpoint tables for interpretation of MICs and zone diameters. [Internet]. Breakpoint tables for interpretation of MICs and zone diameters. 2023 [cited 2023 Jan 25]. Available from: https://www.eucast.org/fileadmin/src/media/PDFs/EUCAST_files/Breakpoint_tables/v_13.0_Breakpoint_Tables.pdf

22. King A, Phillips I, Kaniga K. Comparative *in vitro* activity of telavancin (TD-6424), a rapidly bactericidal, concentration-dependent anti-infective with multiple mechanisms of action against Gram-positive bacteria. Journal of Antimicrobial Chemotherapy. 2004 May;53(5):797–803.

23. Thornsberry C, Draghi D, Benton B, Cohen M, Jones M, Krause K, et al. Baseline profile of telavancin activity against streptococci; results of the USA Surveillance Initiative. In: 46th Annual Interscience Conference on Antimicrobial Agents and Chemotherapy. San Francisco, CA; 2006.

24. Draghi D, Jones M, Flamm R, Thornsberry C, Sahm D. Telavancin activity against current and diverse populations of enterococci and Streptococcus pneumoniae. In: 45th Annual Interscience Conference on Antimicrobial Agents and Chemotherapy. Washington, DC; 2005.

25. Das B, Sarkar C, Das D, Gupta A, Kalra A, Sahni S. Telavancin: a novel semisynthetic lipoglycopeptide agent to counter the challenge of resistant Gram-positive pathogens. Vol. 4, Therapeutic Advances in Infectious Disease. SAGE Publications Ltd; 2017. p. 49–73.

26. Barcia-Macay M, Lemaire S, Mingeot-Leclercq MP, Tulkens PM, van Bambeke F. Evaluation of the extracellular and intracellular activities (human THP-1 macrophages) of telavancin versus vancomycin against methicillin-susceptible, methicillin-resistant, vancomycin-intermediate and vancomycin-resistant Staphylococcus aureus. Journal of Antimicrobial Chemotherapy. 2006 Dec;58(6):1177–84.

27. Gander S, Kinnaird A, Finch R. Telavancin: *In vitro* activity against staphylococci in a biofilm model. Journal of Antimicrobial Chemotherapy. 2005 Aug;56(2):337–43.

28. Goldstein EJC, Citron DM, Merriam CV, Warren YA, Tyrrell KL, Fernandez HT. *In vitro* activities of the new semisynthetic glycopeptide telavancin (TD-6424), vancomycin, daptomycin, linezolid, and four comparator agents against anaerobic gram-positive species and Corynebacterium spp. Antimicrob Agents Chemother. 2004 Jun;48(6):2149–52.

29. Kaniga K, Blosser R, Karlowsky J, Sahm D. *In vitro* activity of telavancin (TD-6424) against Bacillus anthracis. In: 44th Annual Interscience Conference on Antimicrobial Agents and Chemotherapy. Washington, DC; 2004.

30. Asgharian B, Price O, Kabilan S, Jacob RE, Einstein DR, Kuprat AP, et al. Development of a Zealand white rabbit deposition model to study inhalation anthrax. Inhal Toxicol. 2016 Jan 28;28(2):80–8.

31. Cummings JE, Abdo Z, Slayden RA. Improved non-redundant species screening panels for benchmarking the performance of new investigational antibacterial candidates against Category A and B priority pathogens. JAC Antimicrob Resist. 2022 Apr 1;4(2).

32. Reyes N, Skinner R, Benton BM, Krause KM, Shelton J, Obedencio GP, et al. Efficacy of telavancin in a murine model of bacteraemia induced by methicillin-resistant Staphylococcus aureus. Journal of Antimicrobial Chemotherapy. 2006 Aug;58(2):462–5.

33. Madrigal AG, Basuino L, Chambers HF. Efficacy of telavancin in a rabbit model of aortic valve endocarditis due to methicillin-resistant Staphylococcus aureus or vancomycin-intermediate Staphylococcus aureus. Antimicrob Agents Chemother. 2005 Aug;49(8):3163–5.

34. Abdelhady W, Bayer AS, Gonzales R, Li L, Xiong YQ. Telavancin is active against experimental aortic valve endocarditis caused by daptomycin- and methicillin-resistant Staphylococcus aureus strains. Antimicrob Agents Chemother. 2017 Feb 1;61(2).

35. Jame W, Basgut B, Abdi A. Efficacy and safety of novel glycopeptides versus vancomycin for the treatment of gram-positive bacterial infections including methicillin resistant Staphylococcus aureus: A systematic review and meta-analysis. Vol. 16, PLoS ONE. Public Library of Science; 2021.

36. Cummings JE, Abdo Z, Slayden RA. Improved non-redundant species screening panels for benchmarking the performance of new investigational antibacterial candidates against Category A and B priority pathogens [Supplementary Material]. JAC Antimicrobial Resistance. 2022 Mar 24;4(2).

37. Henderson DW, Peacock S, Belton FC. Observations on the prophylaxis of experimental pulmonary anthrax in the monkey. Journal of Hygiene. 1956;54(1):28–36.

38. Heine HS, Bassett J, Miller L, Hartings JM, Ivins BE, Pitt ML, et al. Determination of antibiotic efficacy against Bacillus anthracis in a mouse aerosol challenge model. Antimicrob Agents Chemother. 2007 Apr;51(4):1373–9.

39. Loving CL, Kennett M, Lee GM, Grippe VK, Merkel TJ. Murine aerosol challenge model of anthrax. Infect Immun. 2007 Jun;75(6):2689–98.

40. Bower WA, Schiffer J, Atmar RL, Keitel WA, Friedlander AM, Liu L, et al. Use of Anthrax Vaccine in the United States: Recommendations of the Advisory Committee on Immunization Practices. MMWR Recommendations Report. 2019;68:1–14.

41. Anthrax Preparedness Authors. Anthrax Preparedness. Clinical Infectious Diseases. 2022 Oct 15;75(Supplement 3).

42. Person MK, Cook R, Bradley JS, Hupert N, Bower WA, Hendricks K. Systematic Review of Hospital Treatment Outcomes for Naturally Acquired and Bioterrorism-Related Anthrax, 1880-2018. Vol. 75, Clinical infectious diseases : an official publication of the Infectious Diseases Society of America. NLM (Medline); 2022. p. S392–401.

43. Stucki A, Gerber P, Acosta F, Cottagnoud M, Cottagnoud P. Efficacy of telavancin against penicillin-resistant pneumococci and Staphylococcus aureus in a rabbit meningitis model and determination of kinetic parameters. Antimicrob Agents Chemother. 2006 Feb;50(2):770–3.

44. Sahm D, Benton B, Jones R, Krause K, Thornsberry C, Draghi D. Interaction of telavancin and other antimicrobial agents tested against key gram positive species. In: 46th Annual Interscience Conference on Antimicrobial Agents and Chemotherapy. San Francisco, CA; 2006.

45. National Institutes of Allergy and Infectious Diseases. *In Vitro* Assessment for Antimicrobial Activity Program [Internet]. 2019 [cited 2023 Jan 25]. Available from: https://www.niaid.nih.gov/research/vitro-assessment-antimicrobial-activity

46. CLSI. M100: Performance Standards for Antimicrobial Susceptibility Testing, 32nd Edition. Wayne, PA; 2022 Feb.

47. CLSI. M45: Methods for Antimicrobial Dilution and Disk Susceptibility Testing of Infrequently Isolated or Fastidious Bacteria, 3rd Edition. Wayne, PA; 2015 Oct.

48. Shaw JP, Seroogy J, Kaniga K, Higgins DL, Kitt M, Barriere S. Pharmacokinetics, serum inhibitory and bactericidal activity, and safety of telavancin in healthy subjects. Antimicrob Agents Chemother. 2005 Jan;49(1):195–201.

